# The WYL domain of the PIF1 helicase from the thermophilic bacterium *Thermotoga elfii* is an accessory single-stranded DNA binding module

**DOI:** 10.1101/163188

**Authors:** Nicholas M. Andis, Christopher W. Sausen, Ashna Alladin, Matthew L. Bochman

## Abstract

PIF1 family helicases are conserved from bacteria to man. With the exception of the well-studied yeast PIF1 helicases (*e.g*., ScPif1 and ScRrm3), however, very little is known about how these enzymes help maintain genome stability. Indeed, we lack a basic understanding of the protein domains found N- and C-terminal to the characteristic central PIF1 helicase domain in these proteins. Here, using chimeric constructs, we show that the ScPif1 and ScRrm3 helicase domains are interchangeable and that the N-terminus of ScRrm3 is important for its function *in vivo*. This suggests that PIF1 family helicases evolved functional modules fused to a generic motor domain. To investigate this hypothesis, we characterized the biochemical activities of the PIF1 helicase from the thermophilic bacterium *Thermotoga elfii* (TePif1), which contains a C-terminal WYL domain of unknown function. Like helicases from other thermophiles, recombinant TePif1 was easily prepared, thermostable *in vitro*, and displayed activities similar to its eukaryotic homologs. We also found that the WYL domain was necessary for high-affinity single-stranded DNA (ssDNA) binding and affected both ATPase and helicase activities. Deleting the WYL domain from TePif1 or mutating conserved residues in the predicted ssDNA binding site uncoupled ATPase activity and DNA unwinding, leading to higher rates of ATP hydrolysis but less efficient DNA helicase activity. Our findings suggest that the domains of unknown function found in eukaryotic PIF1 helicases may also confer functional specificity and additional activities to these enzymes, which should be investigated in future work.

## INTRODUCTION

DNA helicases are motor proteins that separate double-stranded DNA (dsDNA) into single-stranded DNA (ssDNA) templates to facilitate cellular processes that are necessary to maintain genome integrity, such as DNA replication, recombination, and repair.*^1^* These enzymes are abundant and conserved throughout evolution, with many being essential for life.*^2^* Indeed, mutations in human DNA helicases are often linked to diseases characterized by genomic instability and a predisposition to cancers.*^3^*

The 5′-3′-directed Superfamily Ib PIF1 helicases exemplify all of these traits.*^4^* Members of this protein family are found in bacteria and eukaryotes, *^4, 5^* and in the case of the fission yeast *Schizosaccharomyces pombe*, the single PIF1 helicase Pfh1 is essential for viability.*^6^* There are also multiple lines of evidence suggesting that human PIF1 (hPIF1) may act as a tumor suppressor, *^7-11^* including the fact that mutation of a conserved residue in the PIF1 family signature sequence*^4^* is linked to breast cancer.*^12^*

The *Saccharomyces cerevisiae* genome encodes two PIF1 family helicases: ScPif1 and ScRrm3. Recently, ScRrm3 was shown to physically associate with Orc5, *^13^* a subunit of the hexameric origin recognition complex (Orc1-6) that demarcates all DNA replication origins in eukaryotes. Orc5 binding requires a short motif in ScRrm3’s large N-terminal domain, which otherwise is of unknown function and is predicted to be natively disordered.*^4^* ScRrm3/Orc5 association modulates DNA replication during replication stress and does not require the ATPase/helicase activity of ScRrm3. Intriguingly, bioinformatics analyses have also recently suggested that a PIF1 helicase/Orc1-6 interaction is also important in amoebae.*^14^* It appears that a gene fusion event occurred in *Dictyostelium fasciculatum* between the genes encoding a PIF1 family helicase and the Orc3 subunit, creating a single protein (Pif1-Orc3). Orc3 and Orc5 interact in *S. cerevisiae*, *^15^* so it is tempting to speculate that the *D. fasciculatum* Pif1-Orc3 and Orc5 proteins recapitulate the important ScRrm3/Orc5 interaction found in yeast. The amoeboid work also found PIF1 helicases with accessory domains containing motifs implicated in ubiquitination (Ubox and CUE-like motifs) and single-stranded nucleic acid binding (RNase H1-like RNA binding and CCHC zinc finger motifs).*^14^*

What other accessory domains are found in PIF1 helicases, and how might these domains impact helicase function or be predictive of novel roles for these enzymes (as with *D. fasciculatum* Pif1-Orc3 and ScRrm3/Orc5 above)? Here, we report that the ScPif1 and ScRrm3 helicase domains are interchangeable, and thus, functional specificity is dictated by the domains of unknown function in these enzymes. We also found that several species of thermophilic bacteria encode PIF1 family helicases that contain C-terminal WYL domains, which are predicted ligand binding modules.*^16^* We show that *Thermotoga elfii* Pif1 (TePif1) is thermostable in solution and displays ATPase, DNA binding, and DNA unwinding activities similar to other PIF1 helicases. Mutational analysis indicated that the WYL domain of TePif1 inhibits ATPase activity but stimulates helicase activity. Further, TePif1 lacking its WYL domain or containing mutations that neutralize its predicted positively charged ligand binding channel had reduced ssDNA binding affinity. The recombinant WYL domain in isolation bound ssDNA, suggesting that it functions as an accessory ssDNA binding interface in the context of the full-length TePif1. Thus, the uncharacterized domains of other PIF1 helicases may contain biochemical activities that are also important for function.

## MATERIALS AND METHODS

### Yeast strains, media, and plasmids

All strains (Table 1) were created by standard methods and are derivatives of the YPH499 genetic background (*MATa ura3-52 lys2-801_amber ade2-101_ochre trp1Δ63 his3Δ200 leu2Δ1*).*^17^* Cells were grown in standard *S. cerevisiae* media at 30°C. Plasmids (Table 2) were constructed by standard molecular biology techniques, and details are available upon request. For the spot dilution assays shown in Figure 1, cells were grown to saturation in selective media, diluted to an optical density at 660 nm of 1.0, and then 10-fold serial dilutions were plated on the indicated media.

**Fig. 1.**
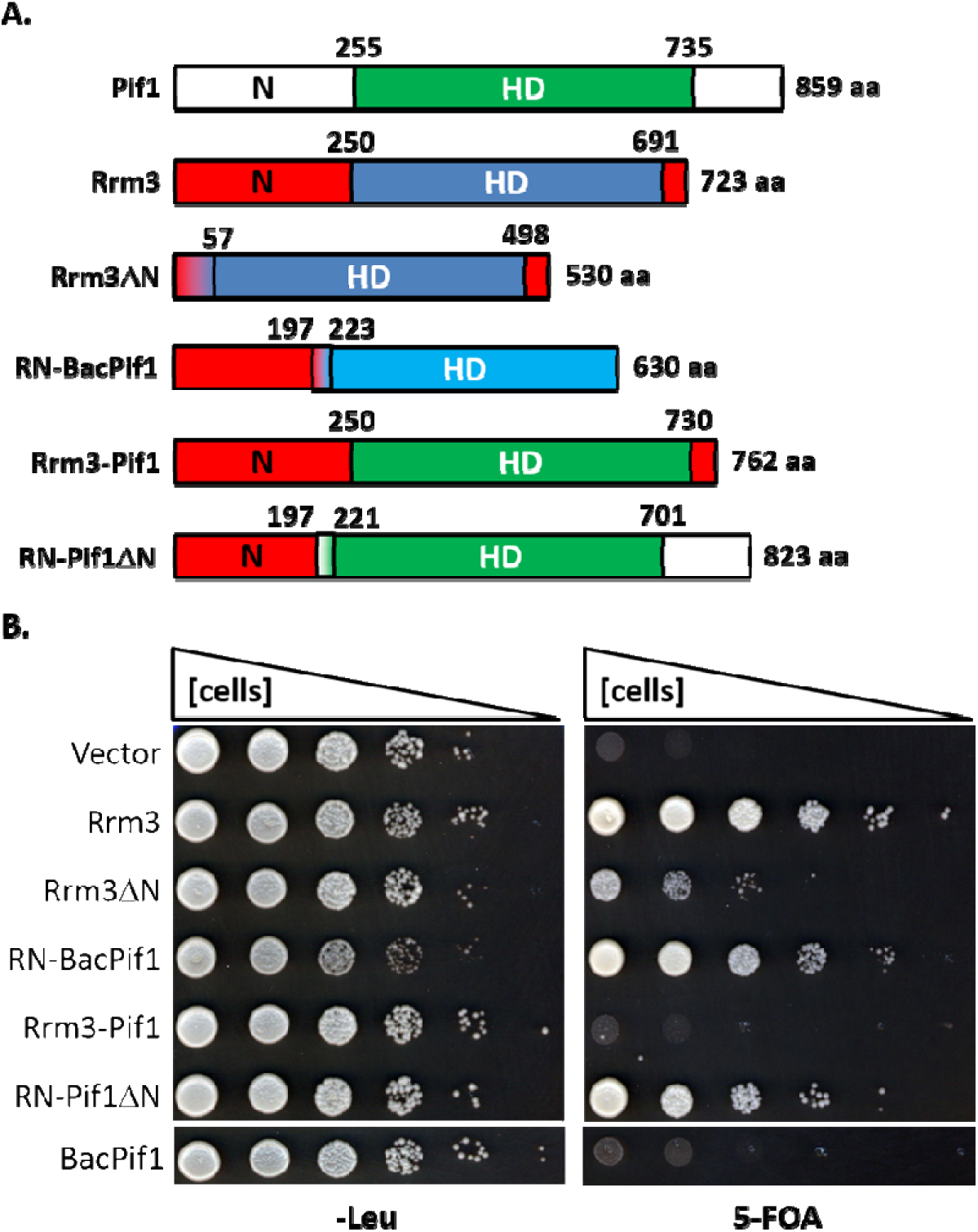
The N-terminus of ScRrm3 is important for function. A) Domain schematics of the helicase constructs tested for rescuing *S. cerevisiae rrm3Δ srs2Δ* synthetic lethality. N denotes the N-terminal domain, and HD denotes the helicase domain. The numbers above the domains indicate the domain boundaries in amino acids. B) Fusing the N-terminus of ScRrm3 to either the helicase domain of ScPif1 (RN-Pif1ΔN) or the Bacteroides sp. 2_1_16 Pif1 protein (RN-BacPif1) yields chimeras capable of rescuing the synthetic lethality of *rrm3Δ srs2Δ* cells. The results shown are representative of three experiments performed with distinct clones of each strain.

**Table 1.**
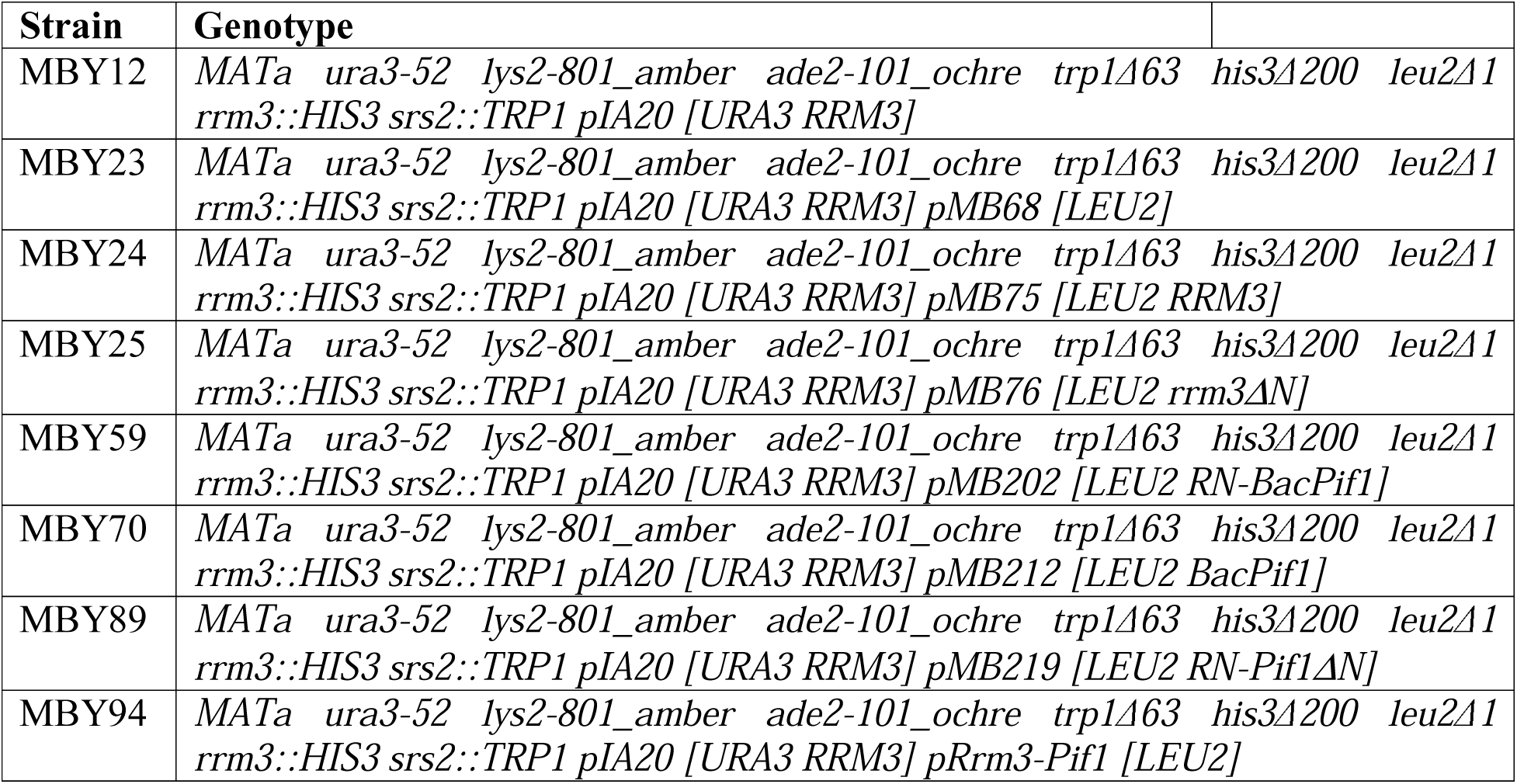
Yeast strains used in this study.

**Table 2.**
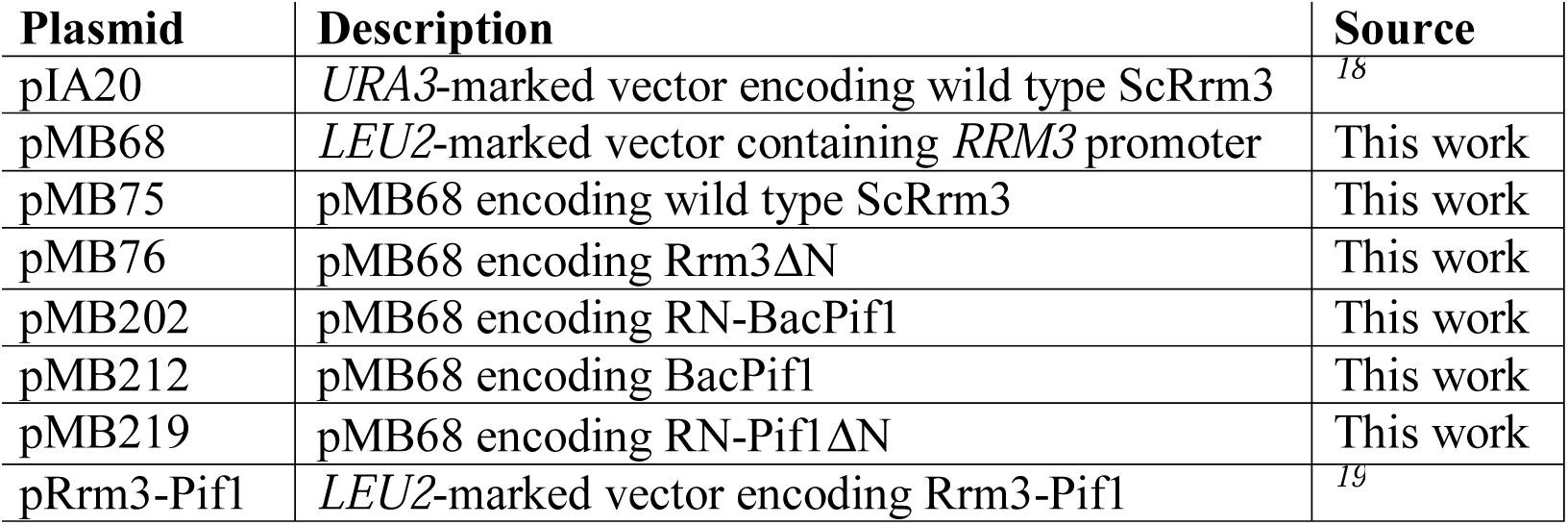
Yeast expression plasmids used in this study.

### Nucleotides, oligonucleotides, and other reagents

ATP was purchased from GE Healthcare (Little Chalfont, UK) or DOT Scientific (Burton, MI, USA). α-^32^P-ATP and γ-^32^P-ATP were purchased from PerkinElmer (Waltham, MA, USA). The oligonucleotides used in this work were synthesized by IDT (Coralville, IA, USA) and are listed in Supplemental Table S1. All enzymes were from New England Biolabs (Ipswich, MA, USA) unless otherwise noted.

### Protein cloning, expression, and purification

A synthetic gBlock DNA encoding the TePif1 protein sequence (Accession no. WP_028843943) was codon optimized for expression in *E. coli* and synthesized by IDT. The DNA was PCR-amplified using oligonucleotides MB1035 and MB1036 (Table S1) and cloned into the *BamHI* and *XhoI* sites of pMB131*^20^* to generate the expression vector pMB427. The sequence of this and all other vectors was verified by DNA sequencing (ACGT, Inc., Wheeling, IL, USA). Details of the TePif1-DENQ, TePif1-ΔWYL, TePif1-4x, WYL, ToPif1, and TyPif1 clonings can be found in the Supplemental Data.

Expression plasmids were transformed into chemically competent Rosetta 2(DE3) pLysS cells and were incubated at 37°C on LB medium supplemented with 100 μg/mL ampicillin and 34 μg/mL chloramphenicol. Fresh transformants were used to inoculate one or more 5-mL LB cultures supplemented with antibiotics and incubated at 30°C for ~6 h with aeration. These starter cultures were then diluted 1:100 in ZYP-5052 autoinduction medium containing 1x trace metals mix *^21^*, 100 μg/mL ampicillin, and 34 μg/mL chloramphenicol and incubated at 22°C with agitation to OD_600_ >3 (15-18 h). Cells were harvested by centrifugation for 10 min at 5,500 x g and 4°C. Cell pellets were weighed and frozen at −80°C prior to lysis or for long-term storage.

Frozen cell pellets were thawed at room temperature by stirring in 4 mL/g cell pellet resuspension buffer (25 mM Na-HEPES (pH 7.5), 5% (v/v) glycerol, 300 mM NaOAc, 5 mM MgOAc, and 0.05% Tween-20) supplemented with 1x protease inhibitor cocktail (Sigma), and 20 μg/mL DNase I. Cells were lysed by six passes through a Cell Cracker operated at >1000 psi. All subsequent steps were performed at 4°C. The soluble fraction was clarified by centrifugation for 30 min at 33,000 x g followed by filtering the supernatant through a 0.22-μm membrane. This mixture was then applied to a Strep-Tactin Sepharose column (IBA) pre-equilibrated in resuspension buffer using an ÄKTA Pure (GE Healthcare Life Sciences). The column was washed with 20 column volumes (CVs) of resuspension buffer, 10 CVs of resuspension buffer supplemented with 5 mM ATP, and 10 CVs of resuspension buffer. Protein was eluted with 15 CVs of resuspension buffer supplemented with 2.5 mM desthiobiotin (IBA). Column fractions were examined on 8% SDS-PAGE gels run at 20 V/cm and stained with Coomassie Brilliant Blue R-250 (BioRad). Peak fractions were pooled, concentrated with Amicon Ultra-4 30K centrifugal filters, and loaded onto a HiPrep 16/60 Sephacryl S-200 HR column (GE Healthcare Life Sciences) pre-equilibrated in gel filtration buffer (25 mM Na-HEPES (pH 7.5), 5% glycerol, 300 mM NaCl, 5 mM MgCl_2_, and 0.05% Tween-20). The protein was eluted with 1.5 CVs gel filtration buffer, and fractions were analyzed by SDS-PAGE as above. Peak fractions were pooled, snap-frozen with liquid nitrogen, and stored at −80°C.

### ATPase assays

ATP hydrolysis was measured via two methods. ATPase activity at various temperatures was analyzed by thin-layer chromatography (TLC) using the method described in *^22^* with some modifications. Briefly, reactions were prepared in helicase buffer (25 mM Na-HEPES [pH 8.0], 5% glycerol, 50 mM NaOAc [pH 7.6], 150 μM NaCl, 7.5 mM MgOAc, and 0.01% Tween-20) and contained 50 nM TePif1, 1 μM oligonucleotide MB551 (Table S1), and 6 μCi of α[^32^P]-ATP. The reactions were incubated for 15 min at temperatures ranging from 4-90°C and stopped by the addition of 2% SDS and 20 mM EDTA. ATP was then separated from ADP by TLC in a buffer composed of 1 M formic acid and 400 mM LiCl for 1 h. The ratio of ATP to ADP was quantified by phosphoimaging using a Typhoon FLA 9500 Variable Mode Imager and ImageQuant 5.2 software. Reactions at each temperature were performed individually and compared to control reactions performed at 20 and 30°C (temperatures empirically determined to yield the highest levels of ATP hydrolysis) at the same time. Background levels of autohydrolysis were subtracted at each temperature tested, and the data were normalized to the highest ATPase activity in each experiment to facilitate comparison of the relative ATPase activity across the temperature range examined.

The ATPase data from assays performed only at 37°C were generated using a NADH-coupled ATPase assay as described in *^23^*. Briefly, reactions were performed in ATPase buffer (25 mM Na-HEPES [pH 8.0], 5% glycerol, 50 mM NaOAc [pH 7.5], 150 μM NaCl and 0.01% NP-40 substitute) supplemented with 5 mM ATP (pH 7.0), 5 mM MgCl_2_, 0.5 mM phospho(enol)pyruvic acid, 0.4 mM NADH, 5 U/mL pyruvate kinase, 8 U/mL lactate dehydrogenase, 50 nM helicase (unless otherwise stated), and 1 μM DNA substrate. Absorbance at 340 nm was read at 37°C in 96-well plates using a BioTek Synergy H1 microplate reader.

Absorbance readings were converted to ATP turnover based on NADH concentration. It was assumed that 1 μM NADH oxidized is proportional to 1 μM ATP hydrolyzed.

### DNA binding assay

DNA binding was measured via gel shifts. The proteins were incubated at the indicated concentrations with 1 nM radiolabeled DNA for 30 min at 37°C in resuspension buffer. Protein–DNA complexes were separated from unbound DNA on 8% 19:1 acrylamide:bis-acrylamide gels in TBE buffer at 10 V/cm. Gels were dried under vacuum and imaged using a Typhoon 9210 Variable Mode Imager. DNA binding was quantified using ImageQuant 5.2 software. All radiolabeled DNA substrates were prepared using T4 polynucleotide kinase and γ-^32^P-ATP by standard methods. See the Supplemental Materials for additional details.

### Helicase assay

DNA unwinding was measured by incubating the indicated concentrations of helicase with 5 mM ATP and 1 nM radiolabeled DNA in resuspension buffer. Reactions were incubated at 37°C for 30 min and stopped with the addition of 1× Stop-Load dye (5% glycerol, 20 mM EDTA, 0.05% SDS and 0.25% bromophenol blue). Unwound DNA was separated on 8% 19:1 acrylamide:bis-acrylamide gels in TBE buffer at 10 V/cm and imaged as for the DNA binding assays. Unwinding time courses were performed under the same conditions but with a single helicase concentration of 50 nM. Reactions were incubated at 37°C for times ranging from 0 to 30 min and stopped as above.

### Statistical analyses

All data were analyzed and plotted using GraphPad Prism 6 (GraphPad Software, Inc). The plotted values are averages, and the error bars were calculated as the standard deviation from three or more independent experiments. Unless otherwise noted, all scatter plots were fit to a hyperbolic curve using the equation Y = (B_max_ · X)/(*k_1/2_* · X), where B_max_ was constrained to 100%. *P*-values were determined by analysis of variance (ANOVA). We defined statistical significance as *p* <0.05.

## RESULTS

### S. cerevisiae PIF1 helicase domains are interchangeable

It was previously shown that the N-terminus of ScRrm3 is important for the function of the helicase *in vivo* and that fusing this domain to the ScPif1 helicase domain cannot rescue the synthetic lethality of a *S. cerevisiae rrm3Δ sml1Δ mec1Δ* mutant.*^19^* These data suggest that both the N-terminal and helicase domains of ScRrm3 endow functional specificity to the enzyme. However, the construction of this Rrm3-Pif1 chimera replaced amino acids immediately N-terminal to the Walker A box of the ScPif1 helicase domain, disrupting a predicted β-sheet (Fig. 1A and S1). To determine if this impacted the *in vivo* function of the Rrm3-Pif1 construct, we created a new chimera that retains the predicted β-sheet and fuses (via a glycine-serine linker) the N-terminus of ScRrm3 to ScPif1 lacking the remainder of its N-terminal domain (RN-Pif1ΔN; Fig. 1A).

We tested the activity of the RN-Pif1ΔN construct in *rrm3Δ srs2Δ* cells, another synthetically lethal double mutant, *^24^* covered by wild-type *RRM3* on a *URA3*-marked plasmid. As shown in Figure 1B, when the cells were forced to lose the *URA3* plasmid by plating on media containing 5-fluoroorotic acid (5-FOA), viability could be maintained by wild-type *RRM3* on a *LEU2*-marked plasmid. However, cells transformed with empty vector were 5-FOA sensitive. As with the *rrm3Δ sml1Δ mec1Δ* cells, *^19^* the Rrm3-Pif1 chimera failed to rescue the *rrm3Δ srs2Δ* synthetic lethality, and cells expressing an ScRrm3 construct lacking the first 193 aa from its N-terminal domain (Rrm3ΔN) were very sick (Fig. 1A,B). However, cells expressing the RN-Pif1ΔN chimera were viable on 5-FOA (Fig. 1B), indicating that fusion of the ScRrm3 N-terminus to the ScPif1 helicase domain in the Rrm3-Pif1 construct disrupted the catalytic activity of the protein. Thus, the helicases domains of ScRrm3 and ScPif1 were interchangeable.

The above data suggest that it is not both the N-terminus and helicase domain of ScRrm3 that confer the specificity of function but, rather, the N-terminal domain alone. To test this hypothesis, we constructed a third chimeric protein comprised of the N-terminus of Rrm3 fused to the *Bacteroides* sp. 2_1_16 PIF1 helicase (RN-BacPif1; Fig. 1A). The *Bacteroides* sp. 2_1_16 PIF1 helicase (BacPif1) alone can substitute for the G-quadruplex maintenance function of ScPif1 when heterologously expressed in *pif1* mutant *S. cerevisiae* cells, *^20^* but it does not rescue the *rrm3Δ srs2Δ* synthetic lethality (Fig. 1B). However, the RN-BacPif1 fusion construct functions nearly as well as wild type ScRrm3 (Fig. 1B). Thus, in this instance, the helicase domain of ScRrm3 is a generic motor, and its N-terminal domain performs an import function in cells lacking the Srs2 helicase.

### Generation of recombinant PIF1 helicases from thermophilic bacteria

What is the essential function of the Rrm3 N-terminus in *srs2Δ* cells? More broadly, what other activities do the non-helicase domains of PIF1 family members possess? We attempted to purify the Rrm3 N-terminus (1-193 aa) to investigate these questions biochemically, but failed to generate a soluble construct. Indeed, a major impediment to working with most eukaryotic PIF1 helicases is the difficulty in over-expressing and purifying sufficient quantities of recombinant protein for biochemistry. For instance, *S. cerevisiae* Pif1 is the most studied PIF1 family helicase in part because the full-length enzyme can be routinely over-expressed and purified from *Escherichia coli.^4^* However, far less is known about *S. cerevisiae* Rrm3 because the full-length protein is exceptionally difficult to purify in an active state.*^25, 26^* To overcome these issues, we and others have previously turned to PIF1 helicases in bacteria, which have proven to be very tractable for *in vitro* experimentation. Indeed, work on G-quadruplex (G4) unwinding *in vitro* and G4 motif maintenance *in vivo* by bacterial PIF1 helicases shows that these activities are conserved across 3 billion years of evolution and not just found in one or more PIF1 helicases from model eukaryotic systems.*^20^* More recently, crystal structures of bacterial PIF1 helicases hold clues to elucidating how ScPif1 can bind to and unwind diverse structures (dsDNA, RNA-DNA hybrids, and G4 DNA).*^27-29^* Using these bacterial helicases is similar to the use of homologous proteins from thermophilic archaea to study difficult-to-purify replication factors from eukaryotes.*^30^* Lacking PIF1 homologs in hyperthermophilic archaea, we instead sought to characterize PIF1 helicases from thermophilic bacteria in an effort to identify enzymes that are robust and amenable to biochemistry and structural biology. By querying the NCBI Protein database with the ScPif1 sequence, we found that multiple thermophilic bacteria encode PIF1 family helicases, including *T. elfii* (TePif1), *Thermus oshimai* (ToPif1), and *Thermodesulfovibrio yellowstonii*

(TyPif1) (Fig. 2A and S2). These proteins contain an N-terminal helicase domain that includes the characteristic PIF1 family signature sequence motif*^4, 5^* (Fig. 2A and B). Additionally, they all include a more C-terminally located UvrD_C_2 domain, which is a domain that adopts a AAA**-**like fold and is found near the C-terminus of many helicases.*^31^*

**Fig. 2.**
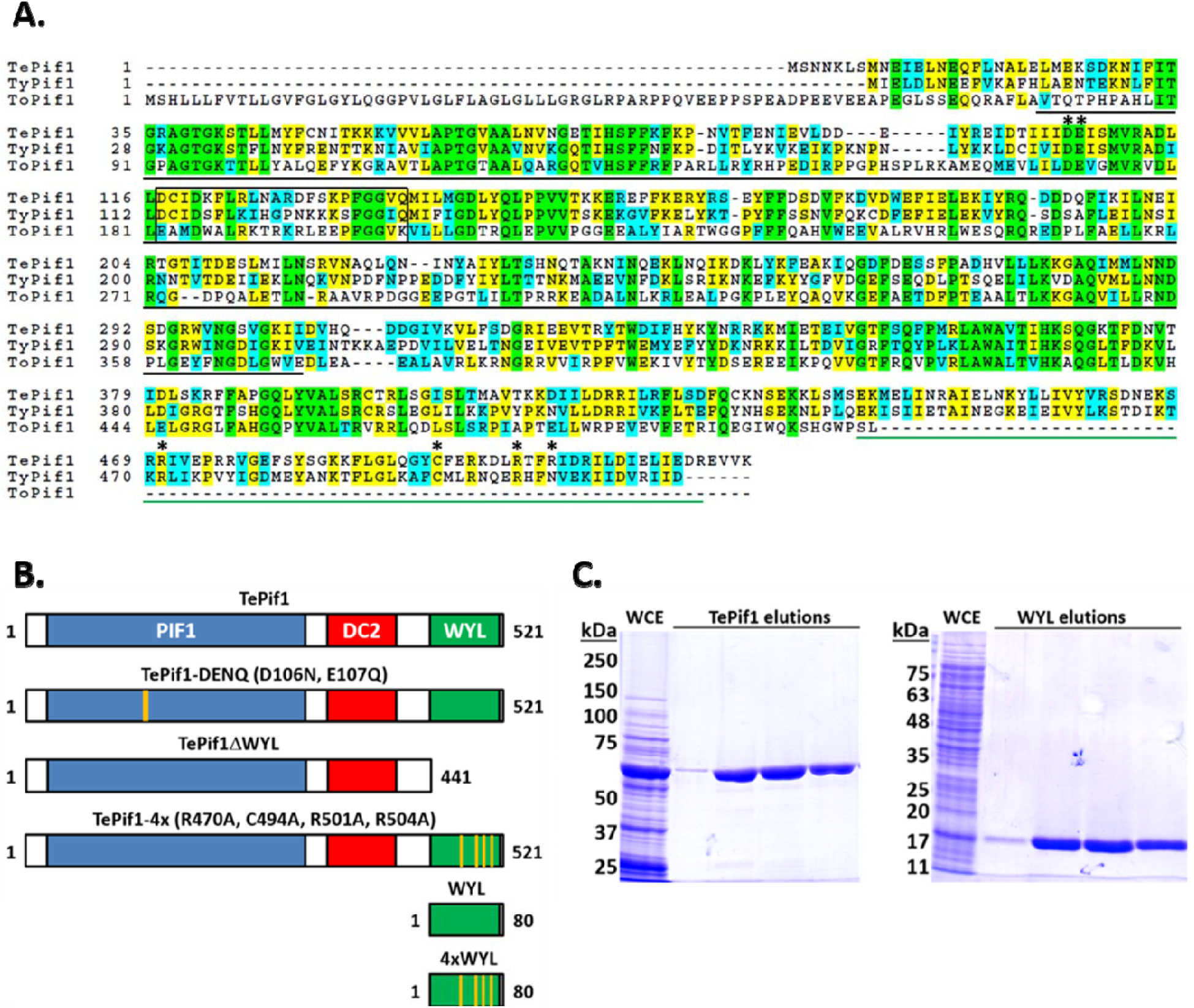
PIF1 family helicases in thermophilic bacteria. A) Sequence alignment of TePif1, TyPif1, and ToPif1. The sequences of the three proteins were aligned using ClustalW, *^32^* and the BOXSHADE program in the Biology WorkBench suite (workbench.sdsc.edu) was used to color-code conserved residues. The completely conserved residues are in green, conserved similarities are in cyan, and identical residues are yellow. The PIF1 family helicase domain is underlined in black, and the PIF1 family signature sequence motif is denoted by the black box. The WYL domain is underlined in green. TePif1 residues that were mutated in this work are marked with asterisks. B) Domain schematics of the proteins that were the focus of this work. The PIF1 family helicase domain (PIF1) is blue, the UvrD_C_2 domain (DC2) is red, and the WYL domain is green. The size of the proteins in amino acids is shown. C) SDS-PAGE of purified recombinant protein preparations. Fractions from the TePif1 (left) and WYL (right) purifications were separated on 8% and 12% acrylamide gels, respectively, and stained with Coomassie blue. The positions of the MW markers are shown.

We also found that a variety of PIF1 helicases from thermophilic bacteria contain a WYL domain at their extreme C-terminus (Fig. 2A and B). Such domains are often found in proteins that interact with nucleic acids, such as the Cas3 protein in type I CRISPR-Cas systems (see *^16^*). In some proteins with tandem WYL domains, it is hypothesized that they may enable oligomerization. Alternatively, based on predicted structural homology with Sm-like SH3 β-barrels and sequence conservation, they may function as binding domains for negatively charged ligands.*^16^* However, very little work has been performed on these proteins. To date, the only characterized WYL domain-containing protein was found to be a negative regulator of one of the CRISPR-Cas systems in *Synechocystis* sp.*^33^*

To begin to characterize PIF1 helicases from thermophilic bacteria and determine the function of the WYL domain, we generated recombinant Te-, To-, and TyPif1 proteins (Fig. 2C and data not shown) by over-expression in *E. coli*. Although all preparations were enzymatically active *in vitro* (Fig. 3-6 and data not shown), wild-type and mutant versions of TePif1 expressed to the highest levels, and thus, TePif1 is the focus of the work herein.

**Fig. 3.**
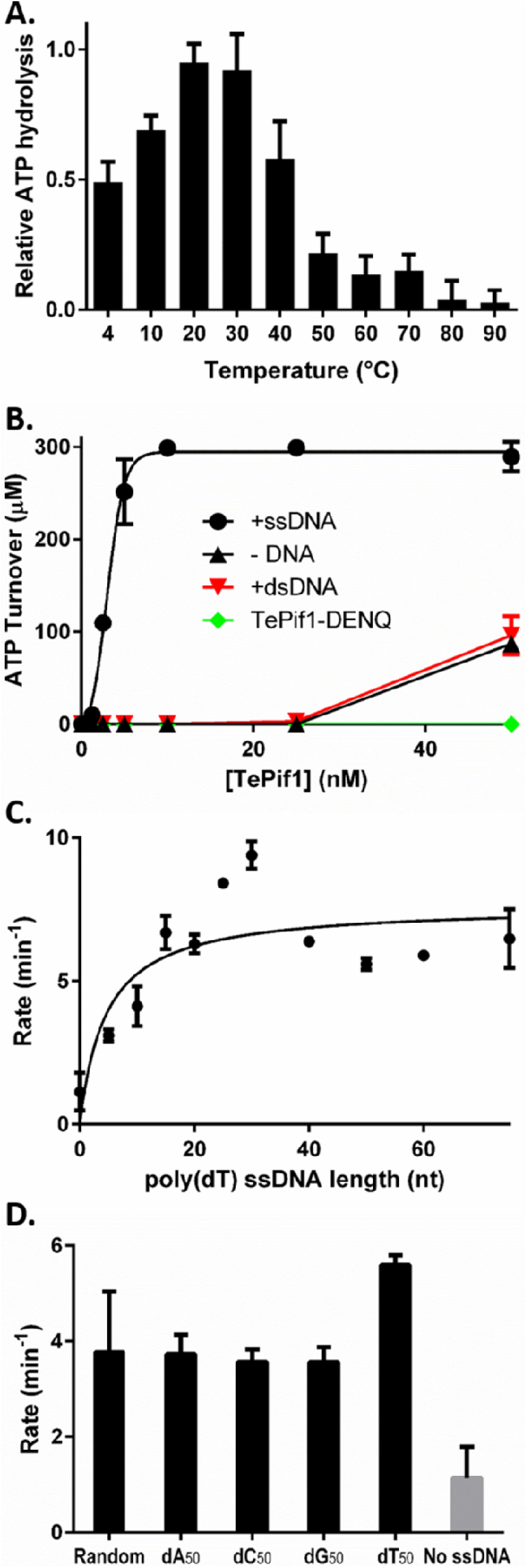
TePif1 ATPase activity is preferentially stimulated by long tracts of poly(dT) ssDNA. A) TePif1 is an active ATPase at a wide range of temperatures. TLC was used to measure TePif1 ATPase activity in the presence of 1 μM poly(dT) 50mer ssDNA at the indicated temperatures. The data are normalized to the highest level of ATP hydrolysis measured in each experiment; see the Methods and Materials for more details. B) TePif1 ATPase activity is greatly stimulated by the presence of ssDNA. The coupled ATPase assay was used to measure ATP hydrolysis by TePif1 at the indicated concentrations in the absence (- DNA) or presence (+DNA) of 1 μM poly(dT) 50mer oligonucleotide. C) The effect of ssDNA length on TePif1 ATPase activity. ATPase hydrolysis by 50 nM TePif1 was measured in the presence of poly(dT) oligonucleotides of the indicated lengths. The slopes (rates) of the hydrolysis curves are plotted *vs*. ssDNA length. D) The effect of ssDNA sequence on TePif1 ATPase activity. The random-sequence, poly(dA), poly(dC), poly(dG), and poly(dT) oligonucleotides were all 50 nt long. In this and all other figures, the data points are averages from ≥ 3 independent experiments, and the error bars represent the standard deviation.

**Fig. 4.**
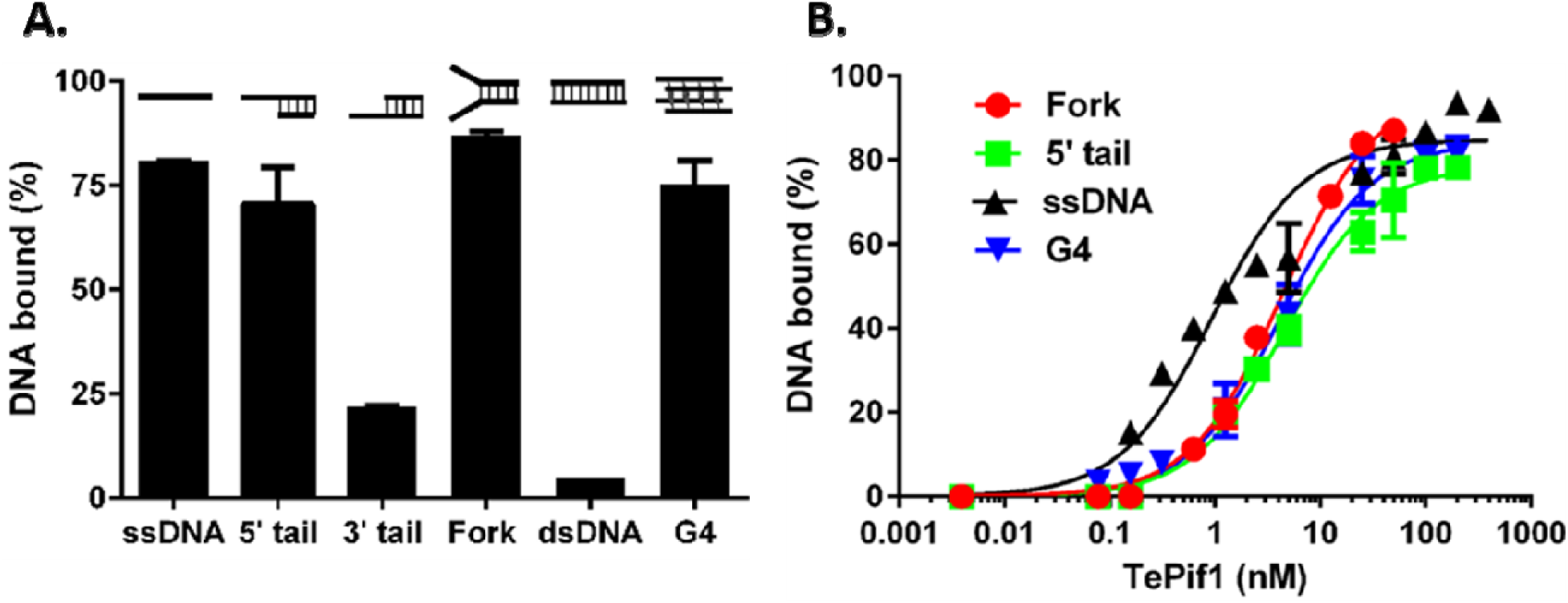
TePif1 preferentially binds ssDNA with high affinity. A) Binding of 50 nM TePif1 to the six substrates shown. The ssDNA is 45-nt long, the ssDNA on the tailed and fork substrates is 25-nt long, and the dsDNA is 25 bp. B) DNA binding as a function of helicase concentration (0-400 nM). TePif1 binds tightly to ssDNA and duplex DNA with 5′ (5′ tail) or 5′ and 3′ ssDNA extensions (Fork). It also tightly binds the TP-G4 DNA substrate (G4). In all cases, the DNA concentration was 1 nM radiolabeled substrate; no unlabeled carrier DNA was used. Representative images of gel shifts are shown in Figure S3.

**Fig. 5.**
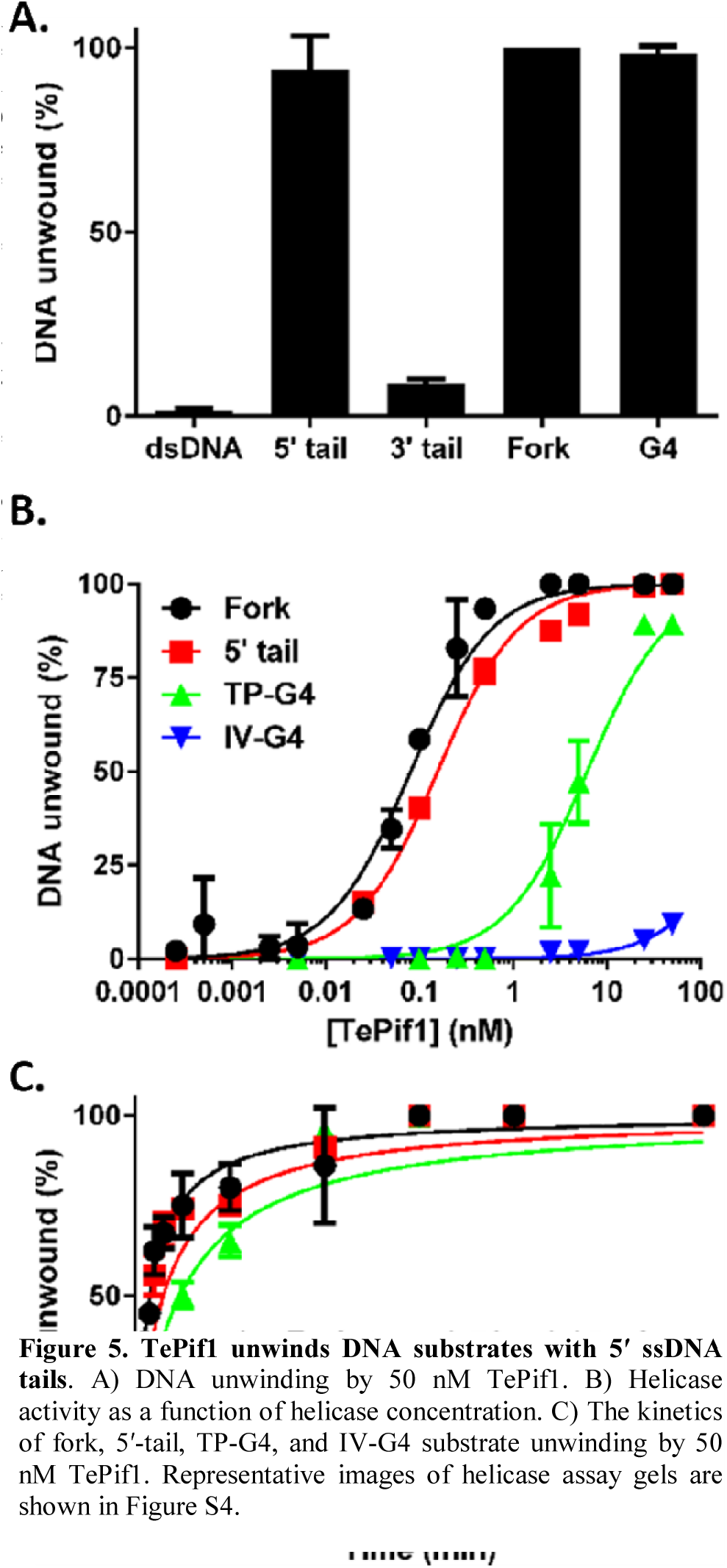
TePif1 unwinds DNA substrates with 5′ ssDNA tails. A) DNA unwinding by 50 nM TePif1. B) Helicase activity as a function of helicase concentration. C) The kinetics of fork, 5′-tail, TP-G4, and IV-G4 substrate unwinding by 50 nM TePif1. Representative images of helicase assay gels are shown in Figure S4.

**Fig. 6.**
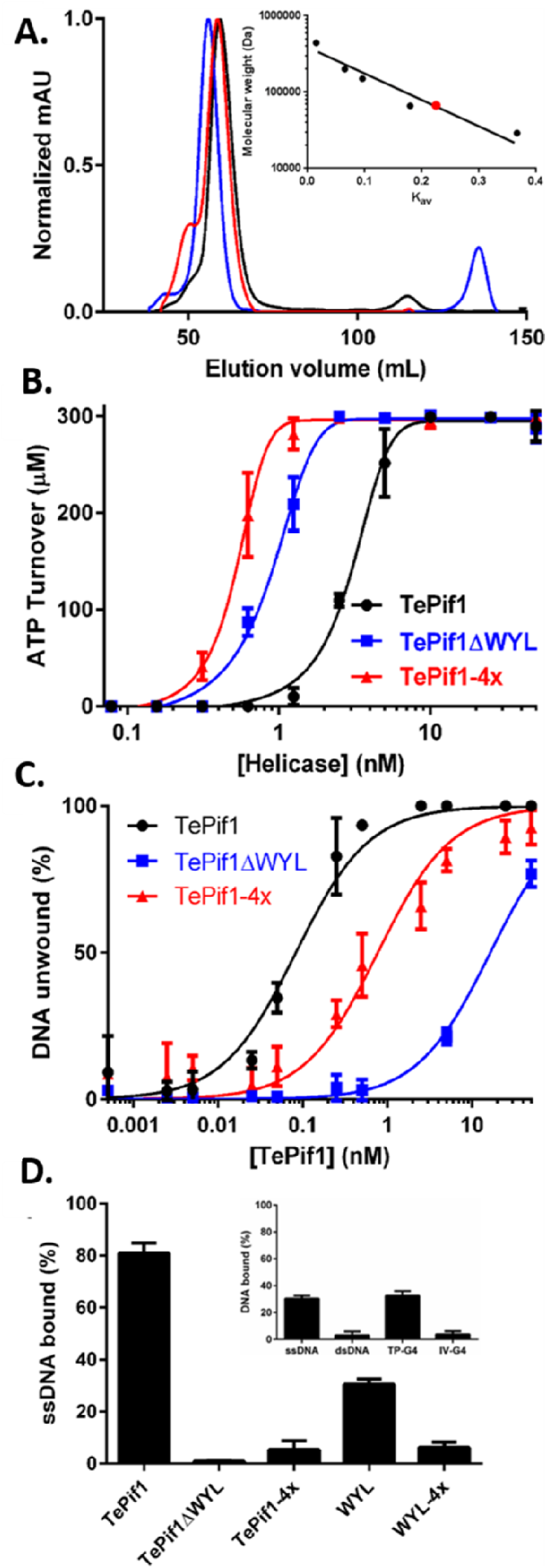
The WYL domain binds ssDNA. A) Gel filtration chromatography trace of wild-type TePif1 and the calculated MW of TePif1 (black), TePif1ΔWYL (blue), and TePif1-4x (red) based on the standard curve (inset). B) TePif1 ATPase activity is stimulated by ablation of the WYL domain. ATP hydrolysis was measured using the coupled assay for the indicated concentrations of TePif1, TePif1ΔWYL, and TePif1-4x. C) Helicase activity on the fork substrate is inhibited by ablation of the WYL domain. D) The TePif1 WYL domain binds ssDNA. Binding by 50 nM wild-type TePif1, truncated protein lacking the WYL domain (TePif1ΔWYL), full-length protein containing R470A, C494A, R501A, and R504A mutations (TePif1-4x), the isolated wild-type WYL domain, and the quadruple R470A, C494A, R501A, and R504A WYL mutant (WYL-4x) is shown. Inset) The WYL domain does not bind to dsDNA or G4 DNA lacking a 5′ ssDNA tail (IV-G4). Binding by 50 nM WYL domain to 1 nM of the indicated substrates is shown. The data shown in all cases is the average from ≥3 independent experiments.

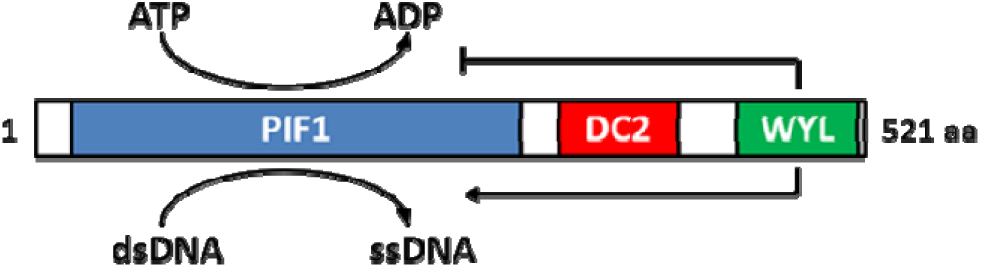

### TePif1 is a DNA-stimulated ATPase

As *T. elfii* grows between 50 and 72°C, with an optimal temperature of 66°C, *^34^* we measured ATP hydrolysis to determine the range of temperatures over which recombinant TePif1 was active *in vitro*. As shown in Figure 3A, significant ATP hydrolysis was observed between 4 and 70°C, with maximum activity at 20-30°C. These data suggest that TePif1 could be a suitable enzyme for use in helicase-dependent amplification (HDA) or other isothermal molecular biology applications. Regardless, because TePif1 was highly active at 37°C, we used this temperature for convenience in the remainder of our biochemical assays.

Like most helicases, the ATPase activi of TePif1 was greatly stimulated by the presen of ssDNA (Fig. 3B, *k_1/2(ATP)_* = 2.96 ± 0.13 nM By including ssDNA substrates of vario lengths, we also found that the stimulation TePif1 ATPase activity was length dependen with a maximal stimulation of ~6-fold in t presence of ≥ 20 nt poly(dT) ssDNA (Fig. 3C However, ssDNA as short as 5 nt was still able stimulate ATPase activity ~2.7-fold, which consistent with ScPif1 being able to bind to ve short lengths of ssDNA.*^35^* In contrast to ssDN dsDNA did not significantly stimulate TePi ATPase activity (Fig. 3B).

Because ScPif1 also preferentially binds to and unwinds G-rich DNA *in vitro*, we assessed the effects of ssDNA sequence on TePif1 activity. Using equimolar concentrations of poly(dA), (dC), (dG), (dT), or random sequence 50mer oligonucleotides, we found that all ssDNAs stimulated TePif1 ATPase activity relative to control reactions lacking ssDNA (all *p* < 0.001 *vs*. no ssDNA) (Fig. 3D). Further, poly(dT) ssDNA preferentially stimulated activity (*p* < 0.01 *vs*. all other ssDNAs) (Fig. 3D). There were no significant differences in the rates of ATP hydrolysis in the presence of the other ssDNAs.

To ensure that the observed ATP hydrolysis was due to TePif1 and not a contaminating enzyme from *E. coli*, we also generated a Walker B box mutant of TePif1 (TePif1-DENQ) by mutating the acidic D106 and E107 residues to their uncharged amide counterparts (D106N, E107Q) (Fig. 3B). The Walker B box is an ATPase motif necessary for hydrolysis, *^36^* and indeed, TePif1-DENQ was unable to hydrolyze ATP (Fig. 3B).

### TePif1 is active on substrates that mimic replication forks and G4 DNA

It is unclear what role(s) bacterial PIF1 helicases play *in vivo*, *^5^* but *in vitro* substrate preference can be indicative of *in vivo* function. For instance, ScPif1 is highly active on G-quadruplex substrates *in vitro* and suppresses genomic instability at sites capable of forming such structures *in vivo*.*^20^* To examine the types of DNA substrates that TePif1 might preferentially act on, we assayed for DNA binding and unwinding of a series of simple substrates *in vitro*. As predicted by the stimulation of ATPase activity in Figure 3, TePif1 bound ssDNA but displayed minimal binding to blunt-ended dsDNA (Fig. 4A, Supplemental Fig. S3). However, substrates containing 5′, 3′, or both ssDNA extensions on duplex DNA were appreciably (> 20%) bound by TePif1. The enzyme preferentially bound ssDNA, 5′-tailed, and fork substrates, as indicated by the high affinity binding (ssDNA *K_d_* = 0.93 ± 0.11 nM, 5′-tail *K_d_* = 4.53 ± 0.39 nM, and fork *K_d_* = 4.40 ± 0.26 nM) to all three substrates (Fig. 4B). There were no significant differences between these three binding affinities (*p* > 0.2). Like ScPif1, *^20^* TePif1 also preferentially bound to G4 DNA (Fig. 4A) formed by a sequence found in the mouse immunoglobulin locus (TP-G4*^37^*). This was also a high affinity substrate (*K_d_* = 4.11 ± 0.41 nM; Fig. 4B).

We next assessed *in vitro* helicase activity. Based on the poor binding to blunt dsDNA (Fig. 4A), it was unsurprising that TePif1 displayed no significant unwinding of this substrate (Fig. 5A). Consistent with the 5′-3′ directionality of PIF1 helicases, *^4^* TePif1 displayed only basal levels of unwinding for the 3′-tailed substrate but was a robust helicase on the 5′-tailed substrate. Similarly, the fork and G4 substrates were unwound to completion (Fig. 5A).

To compare unwinding of the 5′-tail, fork, and TP-G4 substrates, we next assayed for helicase activity as a function of helicase concentration. TePif1 unwound the fork and 5′-tail substrates with similar activity (fork *k_1/2_* = 80 ± 7 pM and 5′-tail *k_1/2_* = 156 ± 9 pM), but the *k_1/2_* for the TP-G4 DNA was much higher at 6.2 ± 6 nM (Fig. 5B). However, the kinetics of unwinding were comparable (*t_1/2_* ≈ 1-2 min) (Fig. 5C). Thus, substrates that resemble replication forks or broken forks retaining 5′ ssDNA are preferred TePif1 substrates *in vitro*. The TP-G4 substrate was also vigorously unwound. Due to its lack of ATPase activity, the TePif1-DENQ mutant was unable to unwind DNA (data not shown).

### The WYL domain impacts ATPase, DNA binding, and helicase activity

We next focused on the role of the TePif1 WYL domain. It is hypothesized that WYL domains may mediate oligomerization, so we sought to determine the oligomeric state of TePif1 in solution. In our preparative gel filtration chromatography, the peak of TePif1 eluted with a calculated molecular weight (MW) of 67.1 kDa (68. 5 kDa MW expected) (Fig. 6A), suggesting that recombinant TePif1 is monomeric. As further evidence that the WYL domain does not mediate oligomerization, recombinant TePif1 lacking this domain (TePif1ΔWYL) or containing mutations (R470A, C494A, R501A, and R504A) in conserved residues in the putative ligand binding channel*^16^*(TePif1-4x; Fig. 2B) also eluted as monomers from gel filtration: TePif1ΔWYL calculated MW = 52.1 kDa, expected MW = 58.8 kDa; and TePif1-4x calculated MW = 63.7 kDa, expected MW = 68.2 kDa (Fig. 6A).

If not oligomerization, then what is the function of the TePif1 WYL domain? To address this question, we compared the biochemical activities of the TePif1ΔWYL and TePif1-4x mutant proteins with wild-type. As shown in Figure 6B, ablation of the WYL domain by deletion or mutation stimulated ATP hydrolysis 3.5-fold and 5.7-fold (respectively) relative to wild-type TePif1 (all in the presence of ssDNA). When these same protein preparations were used in helicase assays, however, DNA unwinding was inhibited by WYL domain ablation (Fig. 6C). The *k_1/2_* for TePif1-4x helicase activity (0.77 ± 0.10 nM) was approximately 10-fold greater than wild-type TePif1, and the apparent *k_1/2_* for TePif1ΔWYL helicase activity (16.4 ± 1.1 nM) was nearly 200-fold greater than wild-type.

Based on our biochemical observations, one factor that affects both ATPase and helicase activity is ssDNA. Thus, we hypothesized that the TePif1 WYL domain may be a ssDNA binding module. When we assessed the ssDNA binding activity of TePif1ΔWYL, it was completely eliminated at 50 nM (Fig. 6D) and only reached levels significantly above background at > 200 nM protein (Supplemental Figure S5). This suggested that ssDNA is a ligand bound by the TePif1 WYL domain.

To further investigate this possibility, we used the TePif1-4x mutant. The R470A, C494A, R501A, and R504A mutations eliminate positive charge in the putative ligand binding channel in the WYL domain, and we predicted that ssDNA binding would thus also be impaired in this mutant. As shown in Figure 6D, this hypothesis proved correct; 50 nM TePif1-4x bound only ~5% of the ssDNA substrate, an approximately 15-fold reduction relative to the wild-type protein.

Finally, we over-expressed and purified the TePif1 WYL domain in isolation (Fig. 2B and C) to test for its ability to bind to ssDNA. In gel shift assays, the WYL domain did bind to a poly(dT) 50mer substrate (Fig. 6D), and mutation of the four residues in the putative binding channel in the WYL domain (4xWYL mutant) reduced ssDNA binding. To assay for substrate specificity, we tested binding to dsDNA and G4 substrates. Like full-length TePif1, the isolated WYL domain failed to bind dsDNA (Fig. 6D, inset) unless it had a 5′ ssDNA tail (data not shown). The WYL domain also bound to the TP-G4 structure, but this substrate contains a 21-nt 5′ ssDNA tail (Table S1). To determine if the WYL domain bound to the G4 structure itself or only the ssDNA portion of the structure, we tested another G4 substrate lacking a 5′ ssDNA tail and containing internal ssDNA loop regions of only 1 and 2 nt (IV-G4). The WYL domain failed to bind IV-G4 DNA above background levels, suggesting that it is specific for ssDNA binding. Together, all of these data correspond to the wild-type TePif1, TePif1ΔWYL, and TePif1-4x results, demonstrating that the TePif1 WYL domain is an accessory ssDNA binding module. The implications of these findings are discussed below.

## DISCUSSION

Eukaryotic PIF1 family helicases, especially those in metazoa, generally contain a centrally located helicase domain flanked by N- and C-terminal domains that are often large and of unknown function.*^4^* In contrast, many bacterial PIF1 helicases are smaller proteins comprised of only the helicase domain.*^4, 5^* However, it has recently been appreciated that PIF1s from a variety of prokaryotic and eukaryotic organisms contain fusions of the PIF1 helicase core to domains with known or predicted enzymatic activities.*^14^* Further, as demonstrated above, the ScRrm3 helicase domain is a generic motor that is connected to a functionally important N-terminal domain (Fig. 1). To better understand the roles of non-helicase domains in PIF1 family proteins, we investigated the function of the C-terminal WYL domain that is found in many PIF1 helicases from thermophilic bacteria. In the context of TePif1, we found that the WYL domain is an accessory ssDNA binding module important for regulating the biochemical activities of the helicase.

### The WYL domain provides another DNA binding interface for TePif1

The data in Figure 6 indicate that the TePif1 WYL domain binds ssDNA. However, this is not the only DNA binding interface in TePif1. Although the TePif1-ΔWYL and TePif1-4x mutant proteins bound ssDNA poorly at a concentration of 50 nM, increasing the helicase concentration to 200-500 nM revealed ssDNA binding of up to ~60% (Fig. S5). Thus, the WYL domain appears to be needed for high affinity ssDNA binding, but other motifs in TePif1 can bind to ssDNA with lower affinity. This could explain why helicase activity decreased for the TePif1ΔWYL and TePif1-4x proteins (Fig. 6C). A lower affinity for the substrate and greater off-rate due to the lack of a high affinity ssDNA binding interface would both decrease TePif1 DNA unwinding. In contrast, ATPase activity is increased in the TePif1ΔWYL and TePif1-4x proteins relative to wild type, suggesting that ATP hydrolysis and productive DNA unwinding were partially uncoupled by these mutations.

It is well established that accessory domains in DNA helicases affect the biochemistry of the main DNA unwinding engines themselves. For instance, the C-terminal domain V of the *Methanothermobacter thermautotrophicus* Hel308 helicase contains a secondary ssDNA binding site.*^38^* Much like the WYL domain mutants of TePif1 (Fig. 6C), mutation of a key arginine residue (R647) in domain V stimulates the ATPase activity of the domain IV ratchet-like helicase motor. In contrast, the accessory ssDNA binding site in the XPD helicase does not affect ATP hydrolysis.*^39-41^* Rather, it is hypothesized that this auxillary ssDNA binding domain is responsible for the frequent back stepping of XPD observed using optical tweezers.*^41^* Single molecule techniques will be necessary to determine if the TePif1 WYL domain affects the processivity of unwinding by a similar mechanism.

Perhaps the TePif1 helicase domain enables binding to dsDNA-ssDNA junctions, while the WYL domain is exclusively involved in interactions with ssDNA. A similar phenomenon has been observed for ScPif1 using single molecule experiments, where the helicase does not translocate along ssDNA but localizes to a dsDNA-ssDNA junction and reiteratively reels in ssDNA.*^42^* ScPif1 lacks a WYL domain, but it does contain N- and C-terminal domains of unknown function*^4^* that could harbor a cryptic ssDNA binding interface.

It should be noted that some helicases that are highly homologous to TePif1, such as ToPif1 (35% identity, 56% similarity), lack a WYL domain (Fig. 2A). Thus, they may display a different spectrum of DNA substrate preferences. In the case of ToPif1, however, its domain organization is slightly different from that of TePif1 and TyPif1, which have well defined helicase, UvrD_C_2, and WYL domains (Fig. 2A). In ToPif1, the UvrD_C_2 domain is larger than in TePif1 and TyPif1, and it is predicted to overlap with the C-terminal portion of the helicase domain. Perhaps this unique domain arrangement generates a ssDNA binding surface that can function in a similar manner to the WYL domain found in TePif1.

### What is the *in vivo* role of TePif1?

To date, it is still unclear what role(s) PIF1 family helicases play in bacteria, even in species like *T. elfii* that encode just a single family member.*^5^* The only indications that we do have come from biochemical data generated using recombinant proteins. Aside from an ability to bind and unwind G-quadruplex DNA structures that is conserved across PIF1 helicases from multiple bacterial phyla*^20^* (Fig. 4 and 5), DNA structures that mimic simple replication forks or broken forks with 5′ ssDNA tails appear to be preferred substrates for TePif1 (Fig. 4 and 5) and the Pif1 helicase from *Bacteroides sp. 3_1_23*.*^28^*

By analogy to the predicted roles of *S. cerevisiae* Rrm3 and *S. pombe* Pfh1 as accessory replicative helicases needed for efficient replication fork progression past nucleoprotein barriers, *^4, 43-47^*. TePif1 and other bacterial PIF1 helicases may serve a similar function during replication. It is unclear which replisome components make direct physical contact with Rrm3 and Pfh1, but the Mcm2-7 replicative helicase*^48^* is a likely candidate. The Mcm2-7 complex is the lynchpin of eukaryotic replication forks as a key platform for protein-protein interactions and as a point of regulation.*^49^* Bacteria such as *T. elfii* encode a DnaB-like replicative helicase instead of an Mcm2-7 complex, but analogous replication machinery and regulation is involved in bacterial replisomes.*^50^* Thus, TePif1 may interact with the DnaB-like helicase during replication *in vivo*.

### TePif1 is active across a wide range of temperatures

*T. elfii* can be cultured at temperatures ranging from 50 to 72°C, but fails to grow at 45 and 75°C.*^34^* Recombinant TePif1 displayed ATPase activity at an even wider range of temperatures (4-70°C), but maximal activity occurred at temperatures (20-30°C; Fig. 3A) well below the optimal growth temperature of *T. elfii*. The reason for this phenomenon is unclear, but it could be due to lack of a partner protein that stabilizes or stimulates TePif1 activity *in vivo*. We could not assay for high-temperature helicase activity because the short DNA substrates used in our assays have a T_m_ of 53.4°C, but we predict that temperature would affect helicase activity in a similar manner to ATP hydrolysis. This broad range of temperatures at which TePif1 is active is similar to the appreciable ATPase activity from 37-65°C of the *Picrophilus torridus* MCM helicase.*^51^ P. torridus* is a thermophilic archaeon that grows between 45 and 65°C, with a growth optimum of 60°C.*^52^* The MCM helicases from the thermophilic *Methanobacterium thermoautotrophicum* and hyperthermophilic *Sulfolobus islandicus* display enzymatic activities at similarly wide ranges of temperature.*^53, 54^*

We also found that TePif1 is stable and remains active for DNA unwinding even when stored at room temperature for at least 1 week (data not shown). This fact and the above data suggest that TePif1 may be a valuable enzyme for isothermal molecular biology applications, such as HDA.*^55^* Ideally, the helicase used in HDA can unwind blunt dsDNA, and though TePif1 largely lacks this ability at 37°C, increasing the reaction temperature slightly increases helicase activity on blunt substrates (Figure S6). The addition of the polymerase in the HDA reaction may also stimulate TePif1 activity, a phenomenon that has been observed with the bacteriophage T7 DNA polymerase and replicative helicase combination.*^56^* However, we failed to observe DNA amplification in HDA assays using TePif1 and commercially available DNA polymerases from mesophilic or thermophilic organisms (data not shown). It should be noted that PIF1 family helicases such as TePif1 can unwind a variety of non-canonical DNA secondary structures (Fig. 5), *^4, 20, 57^* so TePif1 may instead be a valuable accessory helicase in traditional HDA reactions that can be inhibited by such structures. Indeed, our ongoing work focuses on the ability of TePif1 to unwind structured DNA and its performance in HDA.

In conclusion, we biochemically characterized the first PIF1 family helicase from a thermophilic bacterium, demonstrating that TePif1 has similar activities to its better studied bacterial and eukaryotic homologs. Further, we also found that the WYL domain in TePif1 functions to bind ssDNA and regulate ATPase and helicase activity. Our data, combined with similar investigations of PIF1 helicases fused to domains with other enzymatic activities, *^14^* will shed light on the *in vivo* functions of PIF1 family proteins in diverse organisms and may suggest roles for the uncharacterized non-helicase domains of eukaryotic PIF1s.

## ASSOCIATED CONTENT

**Supporting Information**.

The following files are available free of charge.

Supplemental experimental procedures, tables, figures, and references (PDF)

## AUTHOR INFORMATION

### Author Contributions

The manuscript was written through contributions of all authors. All authors have given approval to the final version of the manuscript.

## ACKNOWLEDGMENT

We thank Cody Rogers and David Nickens for carefully reading this manuscript, Garrett Booher, Matan Cohen, and Ankon Paul for contributing early results, and members of the Bochman and van Kessel labs for feedback.

## ABBREVIATIONS

dsDNA: double-stranded DNA
ssDNA: single-stranded DNA
hPIF1: human PIF1
TePif1: *Pseudothermotoga elfii* Pif1
ToPif1: *Thermus oshimai* Pif1
TyPif1: *Thermodesulfovibrio yellowstonii* Pif1
HDA: helicase dependent amplification
MW: molecular weight
TePif1ΔWYL: TePif1 lacking its C-terminal WYL domain
TePif1-4x: TePif1 containing R470A, C494A, R501A, and R504A mutations
CV: column volume
TLC: thin-layer chromatography

